# 12R-HETE acts as an endogenous ligand of Nur77 in intestine and regulates ILC3s plasticity

**DOI:** 10.1101/2023.06.26.546623

**Authors:** Ningning Huang, Ling Ye, Hao Li, Hongkui Wei, Jian Peng

## Abstract

Group 3 innate lymphoid cells (ILC3s), a heterogeneous population, are tissue-resident myeloid cells and have an essential role in bacterial infection. Although the plasticity of NKp46^-^ CCR6^-^ double-negative (DN) ILC3s toward the NKp46^+^ ILC3s is an important process in the development of intestinal immunity, the underlying molecular mechanisms responsible for this process remain poorly understood. Nur77 is an orphan receptor which regulates intestinal ILC3s expansion. However, the impact of Nur77 on the plasticity of intestinal ILC3s remains unclear. Here, we generated *Nur77* null mice and investigated ILC3s expansion. The deficiency of *Nur77* inhibited the mouse small intestinal ILC3s expansion and conversion of NKp46^-^ ILC3s to NKp46^+^ ILC3s. We identified that 12R-HETE derived from arachidonic acid (ARA) in mouse intestine is an endogenous ligand of Nur77 and activates its transcriptional activity. The treatment with 12R-HETE promoted the differentiation of NKp46^-^ ILC3s into NKp46^+^ ILC3s by enhancing the T-bet expression, thereby increased IFN-γ production from NKp46^+^ ILC3s, and reduced the susceptibility to bacterial infection in WT, but not Nur77^-/-^, suckling mice. An integrated analysis of ATAC-seq and Smart RNA-seq showed that *Rflnb*, *Impdh1*, *Map1s*, and *Gtpbp3* might be downstream targeted genes of Nur77 in response to 12R-HETE and mediate the regulation of ILC3s plasticity. In the presence of mycophenolic acid, an inhibitor of IMPDH, 12R-HETE no longer regulated the percentages of RORγt^+^ILC3s and NKp46^+^ILC3s. We conclude that 12R-HETE acts as an endogenous ligand of Nur77, and regulates the ILC3s expansion and plasticity, and in turn, gut homeostasis and pathogen defense.

## INTRODUCTION

Innate lymphoid cells (ILCs) play a critical role in the initial immunity against invading pathogens (Panda and Colonna 2019). Among ILCs, ILC3s constitute the predominant population in the intestine and play an important role in maintaining the integrity of the intestinal barrier, regulating the composition of the gut microbiota, and protecting against intestinal infections (Artis and Spits 2015, Withers and Hepworth 2017). Intestinal ILC3s exhibit a remarkable degree of plasticity, allowing these cells to adapt to different microenvironments and differentiate into subsets with distinct functional properties (Gronke et al 2016, Colonna 2018). Intestinal ILC3s are heterogeneous population. Based on the expression of natural cytotoxic receptor NKp46 and CCR6, ILC3s can be classified into CCR6^+^ ILC3s, NKp46^+^ ILC3s and CCR6^-^ NKp46^-^ double negative (DN) ILC3s (Klose et al 2013). DN ILC3s represent intestinal precursors of the NKp46^+^ ILC3s subset by up-regulating the expression of T-bet and activation of Notch signaling (Klose et al 2013, Rankin et al 2013, Chea et al 2016). Although the differentiation of DN ILC3s into NKp46^+^ ILC3s is an important process in the development of intestinal immunity, the underlying molecular mechanisms of this process remain poorly understood.

Nur77, a member of the NR4A subfamily of nuclear receptors, is a crucial contributor to the development and function of many immune cell types, including T cells and macrophages (Hanna et al 2012, Odagiu et al 2021). Recently, Nur77 has been identified as a potential regulator of intestinal ILC3s expansion (Liu et al 2021). However, the impact of Nur77 on the plasticity of intestinal ILC3s remains unclear. Moreover, Nur77 is an orphan nuclear receptor (Hsu et al 2004). Although PGA_2_, an endogenously derived eicosanoid, has been shown to bind to Nur77, which leads to activation of its transcriptional activity based on in vitro small molecule library screening (Lakshmi et al 2019), it remains to be determined whether PGA_2_ exists in the gut and serves as an endogenous ligand for Nur77 (Lakshmi et al 2019). Therefore, the identification of an endogenous ligand of Nur77 in the intestine is important for investigating the role of Nur77 in regulating intestinal ILC3s development.

The aim of this study is to identify the endogenous ligands of Nur77 and explore their roles in regulating the plasticity of ILC3s via Nur77. Specifically, we have determined the endogenous ligands of Nur77 in the intestine by studying protein-metabolite interactions (PMI). Using the wild-type and Nur77 knockout mice, we have investigated the effect of this ligand on ILC3s plasticity regulation. Finally, we have uncovered the key targeted genes that regulate the ILC3s plasticity upon the endogenous ligand-induced Nur77 activation using Smart-Seq and ATAC-Seq and confirmed the modulator effects of the identified genes. Our research results provide novel insights into the critical molecular mechanisms responsible for the development of intestinal innate immunity, and implications for treating intestinal diseases such as inflammatory bowel disease.

## RESULTS

### Nur77 deficiency impairs the expansion and plasticity of intestinal ILC3s in suckling mice

To investigate the role of Nur77 in intestinal ILC3s expansion, we compared the percentages and absolute numbers of ILC3s in small intestinal lamina propria lymphocytes (SI LPLs) of 2-week-old Nur77^+/+^ and Nur77^-/-^ suckling mice. We showed that the percentages and the absolute numbers of ILC3s in Nur77^-/-^ mice were significantly lower than those in Nur77^+/+^mice (Fig. 1A-C). As NKp46^+^ ILC3s are the most abundant subsets of ILC3s in the small intestine (Melo-Gonzalez and Hepworth 2020), we assumed that Nur77 may affect ILC3s subsets. Thus, we determined the percentages of NKp46^+^ ILC3s and NKp46^-^ ILC3s in the small intestine of Nur77^+/+^ and Nur77^-/-^ mice (Fig. 1D). The results showed that the frequencies and numbers of NKp46^+^ ILC3s in total ILC3s were reduced in Nur77^-/-^ mice (Fig. 1E, F). However, the percentages and numbers of NKp46^-^ ILC3s in Nur77 knockout mice were increased compared with that in Nur77^+/+^ mice (Fig. 1G, H). Interestingly, the reduction of NKp46^+^ ILC3s in Nur77^-/-^ mice could not be attributed to the decreased proliferation as indicated by a comparable level of Ki67^+^ cells in NKp46^+^ ILC3s between Nur77^+/+^ and Nur77^-/-^ mice (Fig. 1I), whereas the percentage of Ki67^+^ cells in NKp46^-^ ILC3s was decreased in Nur77^-/-^ mice without an accumulation of NKp46^-^ ILC3s (Fig. 1J). It has been reported that NKp46^-^ ILC3s contained NKp46^+^ ILC3s precursors under the control of T-bet (Klose et al 2013). We found that compared with that in the wild-type controls (Nur77^+/+^ mice), there was a significant increase in the proportion and absolute number of T-bet^+^ DN ILC3s (Fig.1K, L), as well as a higher mean fluorescence intensity (MFI) of T-bet in DN ILC3s in Nur77^-/-^ mice (Fig.1M). These data indicated that Nur77 regulates the development of intestinal ILC3s and the plasticity of DN ILC3s towards the NKp46^+^ ILC3s in suckling mice.

**Fig. 1.**
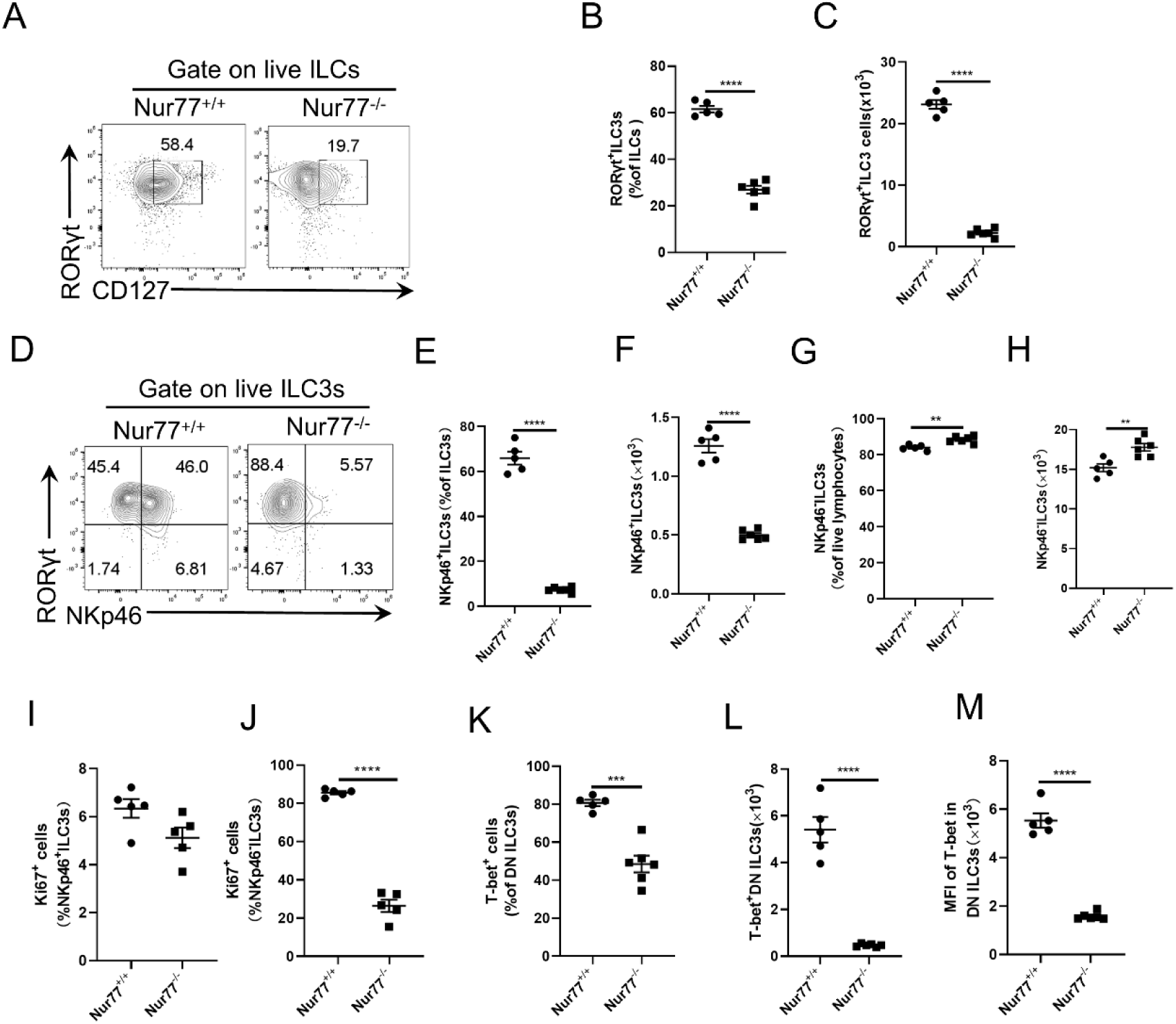
Nur77 deficiency impairs the expansion and plasticity of intestinal ILC3s in suckling mice. The SI LPLs were isolated from *Nur77^+/+^*和*Nur77^-/-^* ^two-week-old mice (A-H). Numbers in flow^ plots represented percentage of RORγt^+^ILC3s on live ILCs gate were shown (A). Flow cytometry was used to detect the percentages of RORγt^+^ILC3s in total ILCs (B). Absolute numbers of RORγt^+^ILC3s (C). Expression of NKp46 gated on total ILC3s was analyzed by flow cytometry (D). Percentages of NKp46^+^ILC3s in ILC3s (E). Absolute numbers of NKp46^+^ILC3s (F). Percentages of NKp46^-^ILC3s in total live lymphocytes (G). Absolute numbers of NKp46^-^ILC3s (H). Flow cytometry was used to detect the percentage of Ki67^+^NKp46^+^ILC3s (I) and Ki67^+^NKp46^-^ILC3s (J). Percentage of Ki67^+^T-bet^+^ cells in DN ILC3s (NKp46^-^CCR6^-^ILC3s) (K). Absolute numbers of T-bet^+^ DN ILC3s (L). MFI of T-bet in DN ILC3s (M). Data are expressed as means ± SEM, n = 5-6. ^a-b^ \**P* < 0.05, \*\**P* < 0.01, \*\*\**P* < 0.001.

### 12R-HETE is an endogenous ligand of Nur77

Previous studies have shown that PGA_2_, a metabolite of ARA, is a potential endogenous ligand of Nur77 (Lakshmi et al 2019). To discover of endogenous ligands for Nur77 in intestine of suckling mice, we applied a metabolomics platform to detect Nur77-LBD metabolite interactions (supplementary Fig. 1A). Here, we used Nur77-LBD to select for small molecules that specifically bind to the ligand-binding pocket of Nur77. Since the natural ligands of Nur77 are lipophilic molecules (Vinayavekhin and Saghatelian 2011, Lakshmi et al 2019), the Nur77-LBD was incubated with lipophilic metabolites extracted from mouse intestine. The eluted sample was analyzed by targeted metabolomic analysis of polyunsaturated fatty acids and their metabolites. We found that the concentration of ARA was higher than docosahexaenoic acid which binding to Nur77-LBD (supplementary Fig. 1B), and enriched 20 small molecules derived from ARA in intestinal extracts. Among them, the most enriched one was 12-HETE, whose level in the eluate of Nur77-LBD was elevated 203.07-fold compared with than in the control. In addition, we also found that PGA_2_ binds to Nur77-LBD in the intestinal extracts, but the relative fold change of PGA_2_ (8.05) was lower than that of 12-HETE (Fig. 2A, B).

**Fig. 2.**
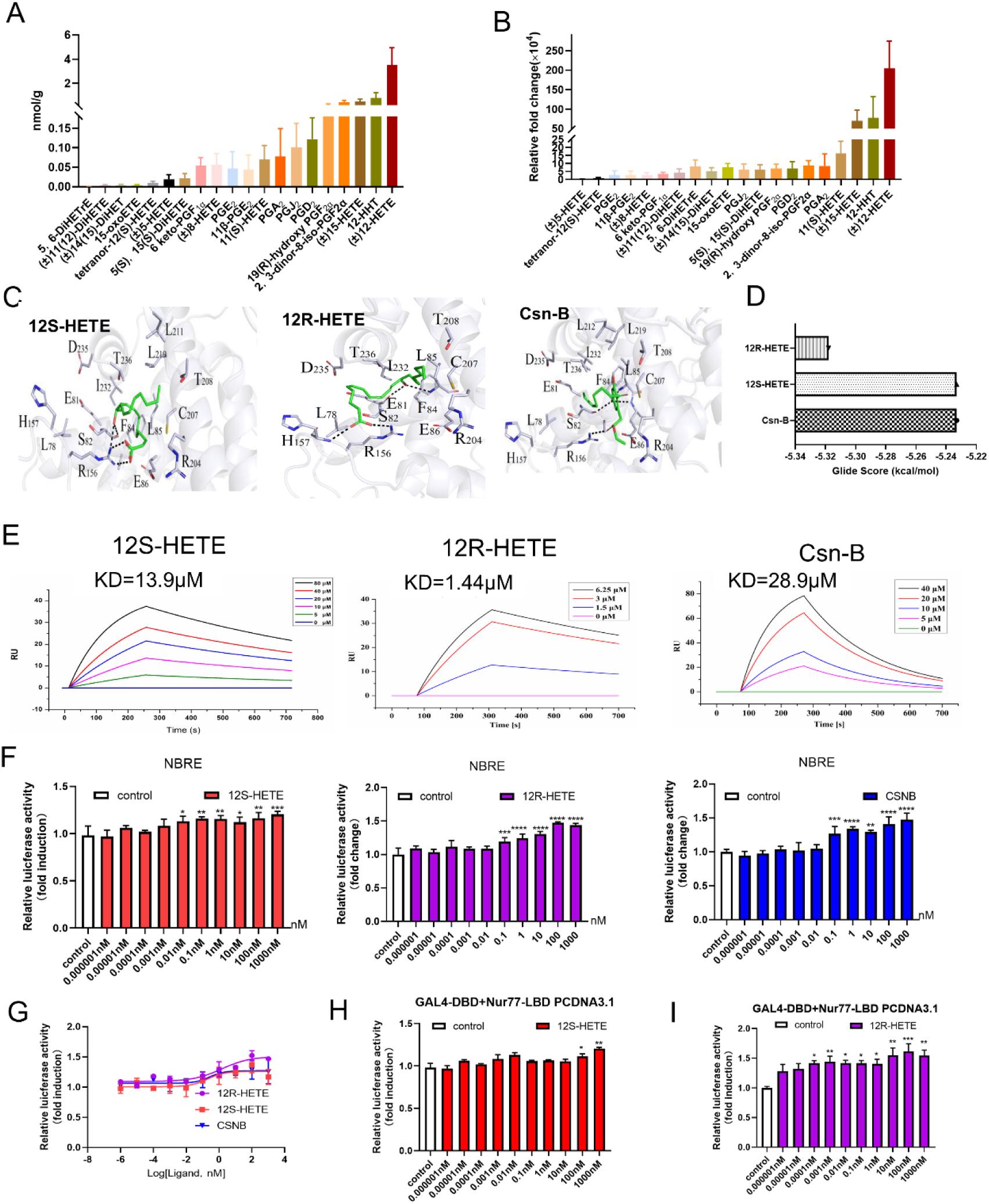
12R-HETE binds to Nur77 and activates its transcriptional activity. Absolute concentration (A) and relative fold change (B) of ARA metabolites bound to Nur77-LBD by LC-MS/MS. The modeling was based on the crystal structure of pig Nur77 (3V3Q) (C-D). Molecular modeling of the pig Nur77 LBD and its interaction respectively with Csn-B, 12S-HETE and 12R-HETE (C) in vitro. Glide Score (D).SPR was used to respectively detect Csn-B、12S-HETE and 12 R -HETE physically binds to Nur77-LBD *in vitro* (E). NBRE and pTK were co-transfected in HEK293T cells, treated with different concentrations of Csn-B, 12S-HETE and 12 R -HETE for 14h, and luciferase activity was examined (F). EC50 graph of Nur77 transcriptional activity activated by Csn-B, 12S-HETE and 12R-HETE, respectively (G). GAL4-DBD, Nur77-LBD and pTK were co-transfected in HEK293T cells, treated with different concentrations of 12R-HETE (H) or 12S-HETE (I) for 14h and luciferase activity was examined.

To confirm the physical binding between 12-HETE and Nur77, molecular docking study was performed to determine the putative docking of R-stereoisomer and S-stereoisomer of 12-HETE. The GLIDE docking scores of 12S-HETE and 12R-HETE were found to be −5.233 and −5.318, respectively. Both of them were comparable to that of an exogenous agonist Csn-B (Fig 2. C, D). By using surface plasmon resonance, we showed that 12R-HETE has a micromolar affinity (KD = 1.44 μmol/L), which is much higher than that of 12S-HETE (KD = 13.9 μmol/L) and CsnB (KD = 28.9 μmol/L) (Fig. 2E). This indicates that 12R-HETE physically interacts with the Nur77-LBD. However, it is unclear whether the Nur77-LBD is required for 12R-HETE to stimulate its transcriptional activity. The results showed that 12R-HETE had an effect on the transactivational activity of Nur77 and the EC_50_ values of 12R-HETE for Nur77 was 14.27×10^-10^mol/L and slightly higher than that of Csn-B (2.627×10^-10^mol/L) (Fig. 2F, G). One previous study reported that GAL4-Nur77 (LBD)-luciferase can be used to screen functional binders for Nur77 (Chintharlapalli et al 2005). Then, HEK293T cells were co-transfected with a chimeric GAL4-Nur77 (full-length Nur77) and GAL4-LBD (ligand binding domain including an AF-2 region with transactivation activity) to investigate the effect of 12R-HETE on the transcriptional activity of Nur77. The treatment with 12R-HETE strongly induced Nur77 transactivational activity, while 12S-HETE was only affected at high concentrations (Fig. 2H, I). These results showed that intestinal 12R-HETE is an endogenous ligand of Nur77 and strongly induces its transactivational activity.

### 12R-HETE promotes intestinal ILC3s expansion and plasticity in a Nur77-dependent manner

As the endogenous ligand of Nur77, 12R-HETE may activate it to induce expansion and plasticity of intestinal ILC3s in suckling mice. To test this hypothesis, we first fed 3d-old mice 12R-HETE until 2-week of age and analyzed the percentages and numbers of ILC3s in lamina propria lymphocytes. We observed that the treatment with 12R-HETE, but not 12S-HETE, increased proportions and numbers of intestinal ILC3s compared to the control (PBS group) (Fig. 3A, B). In addition, higher absolute numbers of IFN-γ^+^ILC3s (Fig. 3C) and enhanced IFN-γ MFI in ILC3s (Fig. 3D, E) were also observed in mice treated with 12R-HETE. There was a trend toward an increase in the proportion of IL-22^+^ ILC3s, while the number of IL-22^+^ ILC3s and IL-22 MFI in ILC3s were increased after 12R-HETE treatment (supplementary Fig. 2A-C). However, 12R-HETE did not affect the percentages and numbers of IL-17A^+^ ILC3s and IL-17A MFI in ILC3s (supplementary Fig. 2D-F). Since we observed that Nur77 knockout impaired plasticity of small intestinal ILC3s and NKp46^-^ ILC3s differentiated into NKp46^+^ ILC3s, we assayed percentages and numbers of NKp46^+^ ILC3s and found that both of them were increased after 12R-HETE administration (Fig. 3F). However, no difference in percentages and numbers of NKp46^-^ ILC3s were found in mice treated with 12R-HETE (Fig. 3G). Moreover, we also showed that 12R-HETE improved the percentages and numbers of IFN-γ^+^ cells in NKp46^+^ ILC3s (Fig. 3H), as well as the higher MFI of IFN-γ in NKp46^+^ ILC3s (Fig. 3I, J). In contrast, neither cytokine production nor plasticity of NKp46^+^ ILC3s was altered after administration of 12S-HETE.

**Fig. 3.**
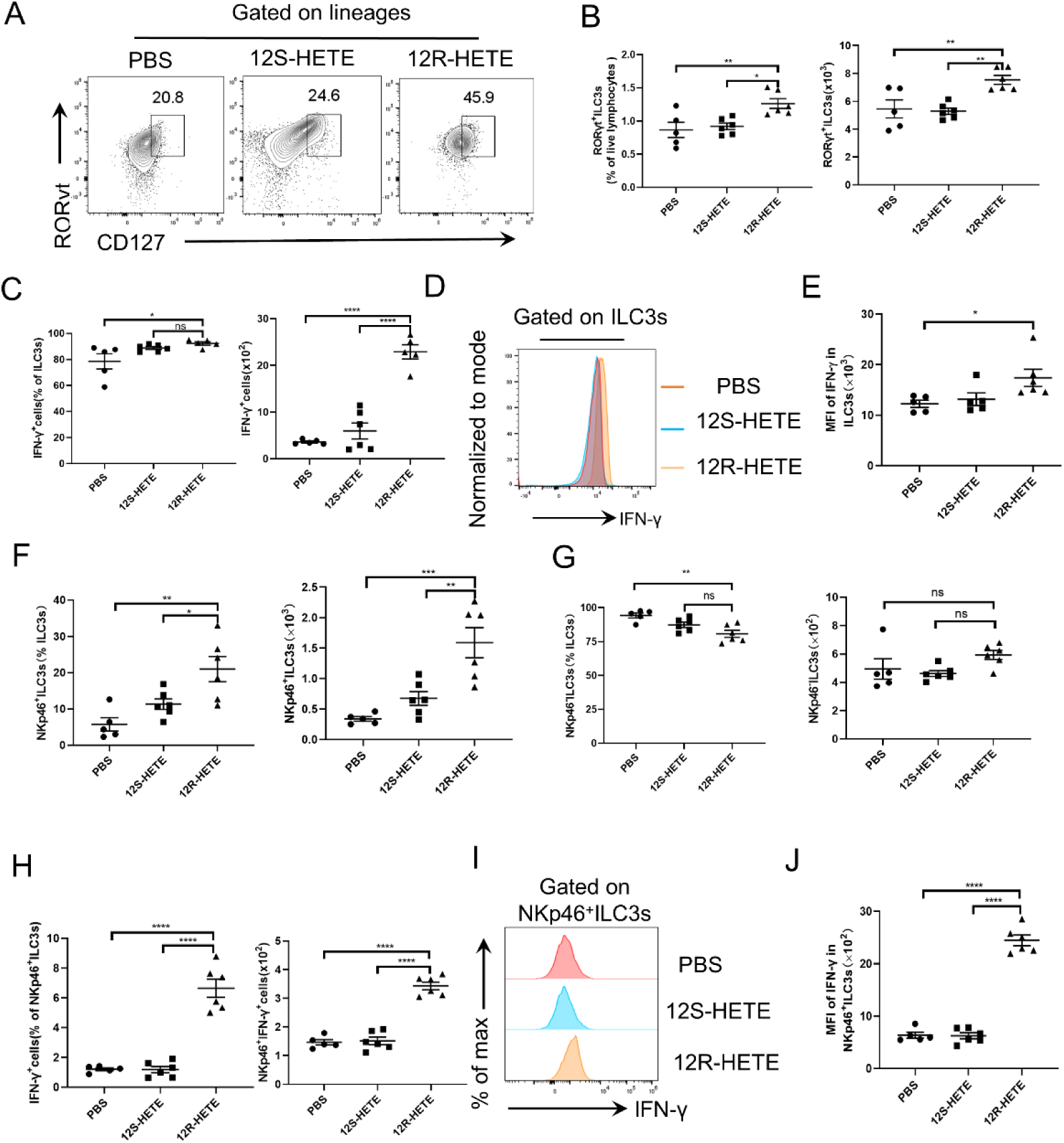
12R-HETE promotes the expansion of ILC3s and IFN-γ production in the small intestine. Mice were given PBS, 1 μmol 12S-HETE and 1 μmol 12R-HETE by gavage for 10 days. Mice were sacrificed at 14 days old, and SI LPLs were extracted. Numbers in flow plots represent percentage of CD127^+^ RORγt^+^ ILC3s on lineage negative cells gate were shown (A). Percentages (left) and absolute numbers (right) of ILC3s in the live lymphocytes from different treatment groups were shown (B). Percentages and cell numbers of IFN-γ from ILC3s in ILC3s from different treatment groups were analyzed (C). Representative histograms show the expression of IFN-γ in ILC3s. The PBS group is red curves, 12S-HETE group is blue curves, and 12R-HETE is orange curves(D). Mean fluorescence intensity (MFI) of IFN-γ in ILC3s (E). Percentages (left) and cell numbers (right) of NKp46^+^ILC3s in ILC3s from different treatment groups were analyzed by flow cytometry (F). Percentages (left) and cell numbers (right) of NKp46^-^ILC3s in ILC3s (G). The percentages of IFN-γ from NKp46^+^ILC3s in ILC3s (left). The numbers of IFN-γ deriving (right) (H). Representative histograms showed IFN-γ expression. The PBS was the orange curves, 12S-HETE was blue curves and 12R-HETE was red curves (I). Mean MFI of IFN-γ in NKp46^+^ILC3s (J). Data are mean ± SEM, n=5-6. Each point represents a biological replicate. \**P* < 0.05, \*\**P* < 0.01, \*\*\*\**P* < 0.0001, ns indicates no significant difference.

The results showed that the increase of NKp46^+^ ILC3s in 12R-HETE group was less likely to be due to decreased apoptotic cells, which was comparable to that in PBS group (supplementary Fig. 2G-I). We observed that 12R-HETE increased the level of Ki67^+^ cells in ILC3s (Fig. 4A, B). A higher proliferation rate of NKp46^-^ ILC3s rather than NKp46^+^ ILC3s was observed in mice treated with 12R-HETE (Fig. 4C, D). Meanwhile, NKp46^+^ ILC3s and DN ILC3s had enhanced T-bet expression after 12R-HETE treatment (Fig. 4E-G).

**Fig. 4.**
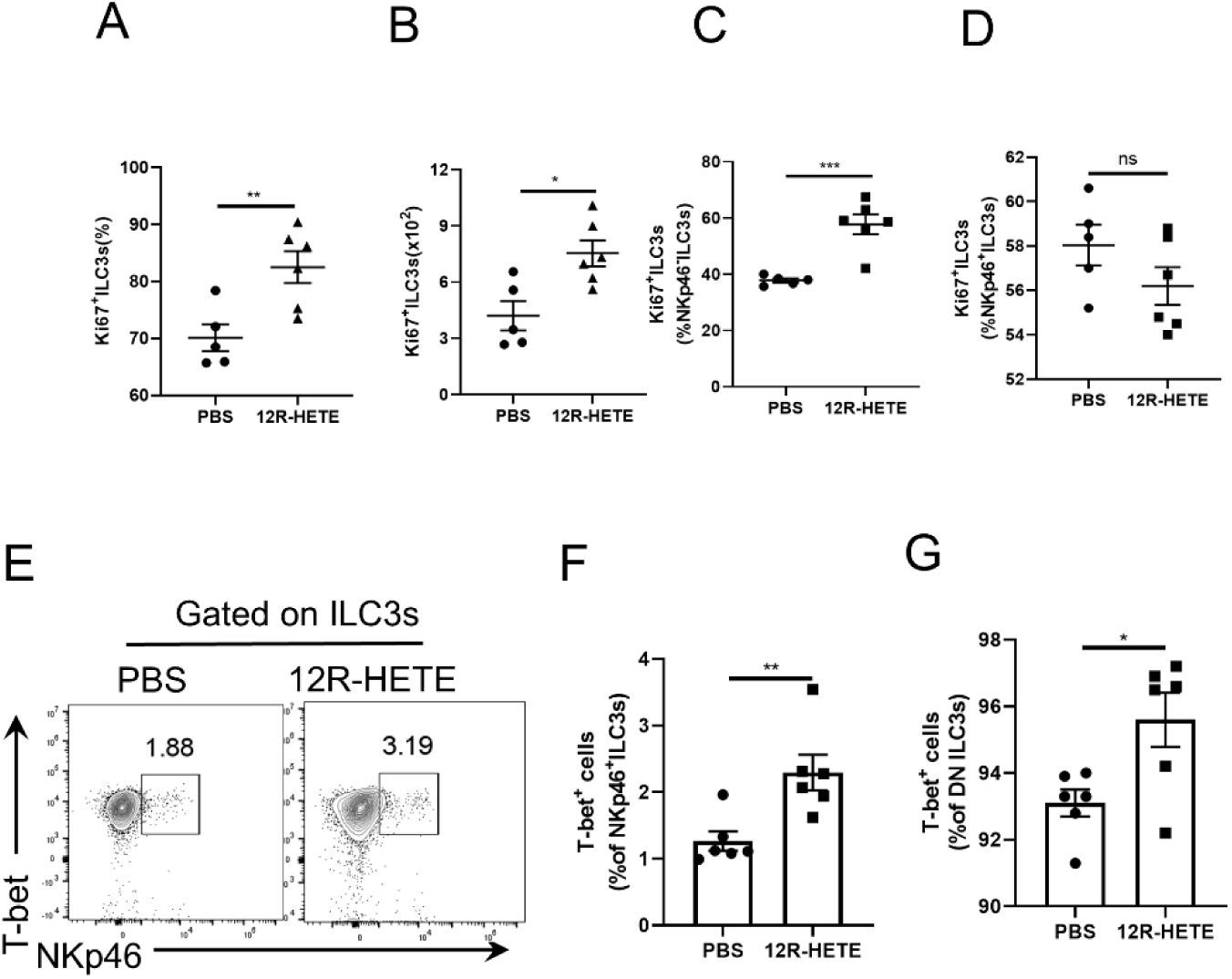
12R-HETE promotes the proliferation of NKp46^-^ILC3s and rapidly differentiates into NKp46^+^ILC3s. Pups were administered with PBS and 1 μmol 12R-HETE for 10 days at five-day-old, respectively. SI LPLs were isolated from mice by flow cytometry. Numbers in flow plots represent percentage of Ki67^+^cells on ILC3s gate were shown (A). Percentage and of Ki67^+^cells in RORγt^+^ILC3s were shown on the left. Absolution numbers of Ki67^+^cells in RORγt^+^ILC3s were shown on the right (B). Percentage of Ki67^+^cells in NKp46^+^ILC3s in intestinal lamina propria (C). Percentage of Ki67^+^cells in NKp46^-^ILC3s (D). Numbers in flow plots represented percentage of T-bet^+^NKp46^+^cells on ILC3s gate were shown (E). Percentage of T-bet^+^cells in NKp46^+^ILC3s (F). Percentage of T-bet^+^cells in DN ILC3s (G). Data show mean ±SEM, n=5-6. \**P* < 0.05, \*\**P* < 0.01, \*\*\**P* < 0.001.

To further confirm whether the 12R-HETE-induced intestinal ILC3s expansion and plasticity depended on Nur77, we fed the Nur77^-/-^ suckling mice with 12R-HETE and then evaluated ILC3s expansion and plasticity again. As expected, the increased percentages of RORγt^+^ILC3s among lymphocytes (Fig. 5A, B), or proportions of NKp46^+^ ILC3s among total ILC3s (Fig. 5D, E) induced by 12R-HETE administration was abolished in Nur77-deficient mice. Moreover, Nur77 deficiency prevented 12R-HETE-induced IFN-γ expression in ILC3s (Fig. 5C) or NKp46^+^ ILC3s (Fig. 5F). These results suggested that 12R-HETE promoted the expansion and plasticity of intestinal ILC3s in a Nur77-dependent manner.

**Fig. 5.**
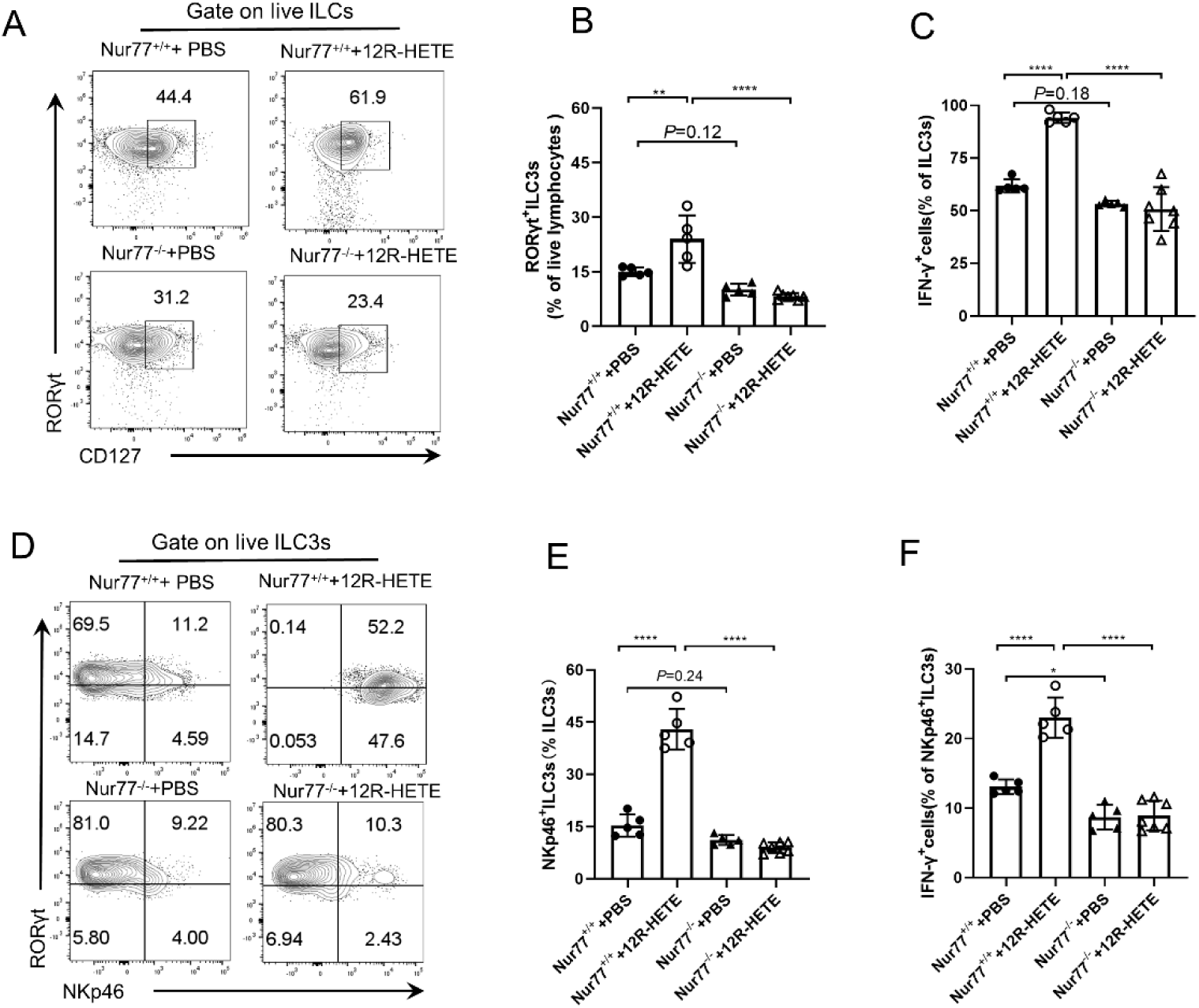
12R-HETE promotes NKp46^+^ILC3s development and IFN-γ production in the small intestine in a Nur77-dependent manner. SI LPLs were isolated from Nur77^+/+^ and Nur77^-/-^ mice at 2-week-old. Numbers in flow plots represent percentages of ILC3s on live lymphocytes gate were shown (A). Flow cytometry assay of the percentages of RORγ^+^ILC3s (B). Percentages of IFN-γ production in ILC3s (C). Numbers in flow plots represent percentages of NKp46^+^ILC3s on ILC3s gate were shown (D). Percentage of NKp46^+^ILC3s in ILC3s (E). Expression of IFN-γ in NKp46^+^ILC3s (F). Data show mean ±SEM, n=5-6. \**P* < 0.05, \*\**P* < 0.01, \*\*\**P* < 0.001.

### 12R-HETE alters the susceptibility to *Salmonella typhimurium*-induced enteritis

NKp46^+^ ILC3s play a key role in host defense of *S*. *typhimurium* infections by producing IFN-γ (Brasseit et al 2018, Kästele et al 2021). We investigated whether 12R-HETE-induced NKp46^+^ ILC3s in intestine of suckling mice alter susceptibility to *S. typhimurium*-induced enteritis. The 12R-HETE-treated mice had increased survival rate, reduced body weight loss, and alleviated intestinal damages compared with PBS-treated controls (Fig. 6 A-C). At d 3 after infection, the *S. typhimurium* load in liver and spleen was decreased in mice pre-treated with 12R-HETE compared to those pre-treated with PBS (Fig. 6 D). Given that 12R-HETE improved ILC3s expansion and regulated ILC3s plasticity toward the NKp46^+^ ILC3s at steady state, we further ask whether those issues exist in the infection state. To address the question, we determined the percentages and numbers of NKp46^+^ ILC3s at 3 d after *S. typhimurium* infection in mice pre-treated with 12R-HETE or PBS. In consistent with the steady-state results, 12R-HETE also improved percentages and numbers of NKp46^+^ ILC3s (Fig. 6 E, F) and MFI of NKp46 in NKp46^+^ ILC3s (Fig. 6 G, H). Additionally, at 3d after infection, the flow cytometry data analysis revealed that 12R-HETE up-regulated the percentages and absolute numbers of IFN-γ^+^ ILC3s (Fig. 6 I-K), together with the increase of IFN-γ MFI in NKp46^+^ ILC3s (Fig. 6 L, M). These data clearly show that 12R-HETE alleviated *S. typhimurium*-induced enteritis through promoting the production of IFN-γ.

**Fig. 6.**
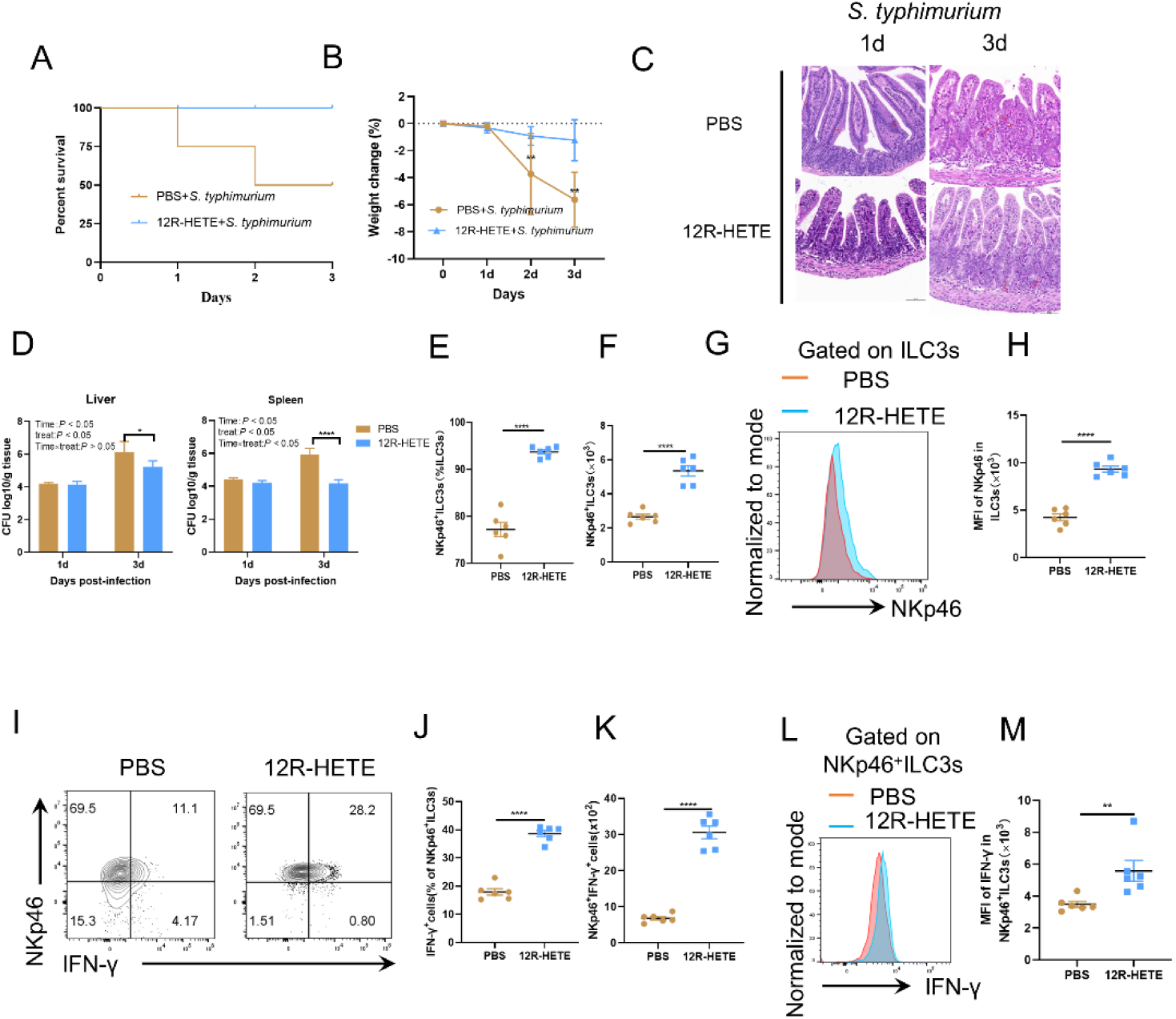
12R-HETE alters the susceptibility to *Salmonella typhimurium*-induced enteritis. Pups were given with PBS and 1μmol 12R-HETE at five-day-old, respectively. 15-day-old mice were administered *S. typhimurium* to infection, mice were slaughter and sampled on days 16 and 18. Survival percentage of mice at1 and 3 days after *S. typhimurium* infection were calculated (A). Weight change (B). *S. typhimurium* in liver and spleen (C). Pathological section of colon stained with H&E (D). Numbers in flow plots represent percentages of RORγt^+^ILC3s on lymphocytes gate were shown on the left. Flow cytometry was used to analyze the percentages and numbers of ILC3s in live lymphocytes were shown (E). Percentages (E) and absolute numbers (F) of NKp46^+^ILC3s in ILC3s were calculated and shown. Representative histograms showed NKp46 expression. The PBS was the orange curves, 12S-HETE was blue curves and 12R-HETE was red curves (G). MFI of NKp46 in NKp46^+^ILC3s (H). Numbers in flow plots represent percentage of IFN-γ expression in NKp46^+^ILC3s on ILC3s gate were shown (I). Percentage of IFN-γ derived cells in NKp46^+^ILC3s were calculated and shown (J). Absolute numbers IFN-γ from cells in NKp46^+^ILC3s (K). Representative histograms showed IFN-γ expression in NKp46^+^ILC3s (L). The expression of IFN-γ in NKp46^+^ILC3s (M). Data show mean ±SEM, n=6-8. \**P* < 0.05, \*\**P* < 0.01, \*\*\**P* < 0.001.

### 12R-HETE promoted the development of NKp46^+^ILC3s by regulation of Nur77 target gene

To elucidate the mechanism by which 12R-HETE activates Nur77 and regulates the plasticity of ILC3s, we sorted small intestinal ILC3s in wild-type mice treated with PBS or 12R-HETE and small intestinal ILC3s in Nur77^-/-^ mice treated with PBS. We then performed assay for transposase-accessible chromatin using high throughput sequencing (ATAC-seq and Smart RNA-seq). Compared to Nur77^+/+^ mice treated with PBS, Nur77^-/-^ mice treated with PBS showed a reduction of 107528 peaks. Conversely, Nur77^+/+^ mice treated with 12R-HETE showed an increase of 52116 peaks compared to Nur77^+/+^ mice treated with PBS. Both datasets shared 39339 peaks (Fig. 7 A), mainly distributed within gene introns and intergenic regions (Fig. 7B). Smart RNA-seq data analysis revealed that Nur77^-/-^ mice treated with PBS showed 5991 upregulated genes and 4378 downregulated genes compared to Nur77^+/+^ mice treated with PBS (supplementary Fig. 3A), whereas Nur77^+/+^ mice treated with 12R-HETE showed 3100 upregulated genes and 4090 downregulated genes compared to Nur77^+/+^ mice treated with PBS (supplementary Fig. 3B). Among these, 296 genes were downregulated when Nur77 was knocked out but upregulated with 12R-HETE treatment (supplementary Fig. 3C), while 430 genes were upregulated when Nur77 was knocked out but downregulated with 12R-HETE treatment (supplementary Fig. 3D). We performed KEGG analysis on the co-upregulated and co-downregulated DEGs and found that these genes were mainly enriched in biological functions such as Notch signaling pathway, AMPK signaling pathway, MAPK signaling pathway and Apoptosis (supplementary Fig. 3E, F). This indicates that Nur77, as a transcription factor, is broadly involved in the regulation of various signaling pathways through the regulation of downstream target gene expression.

**Fig.7.**
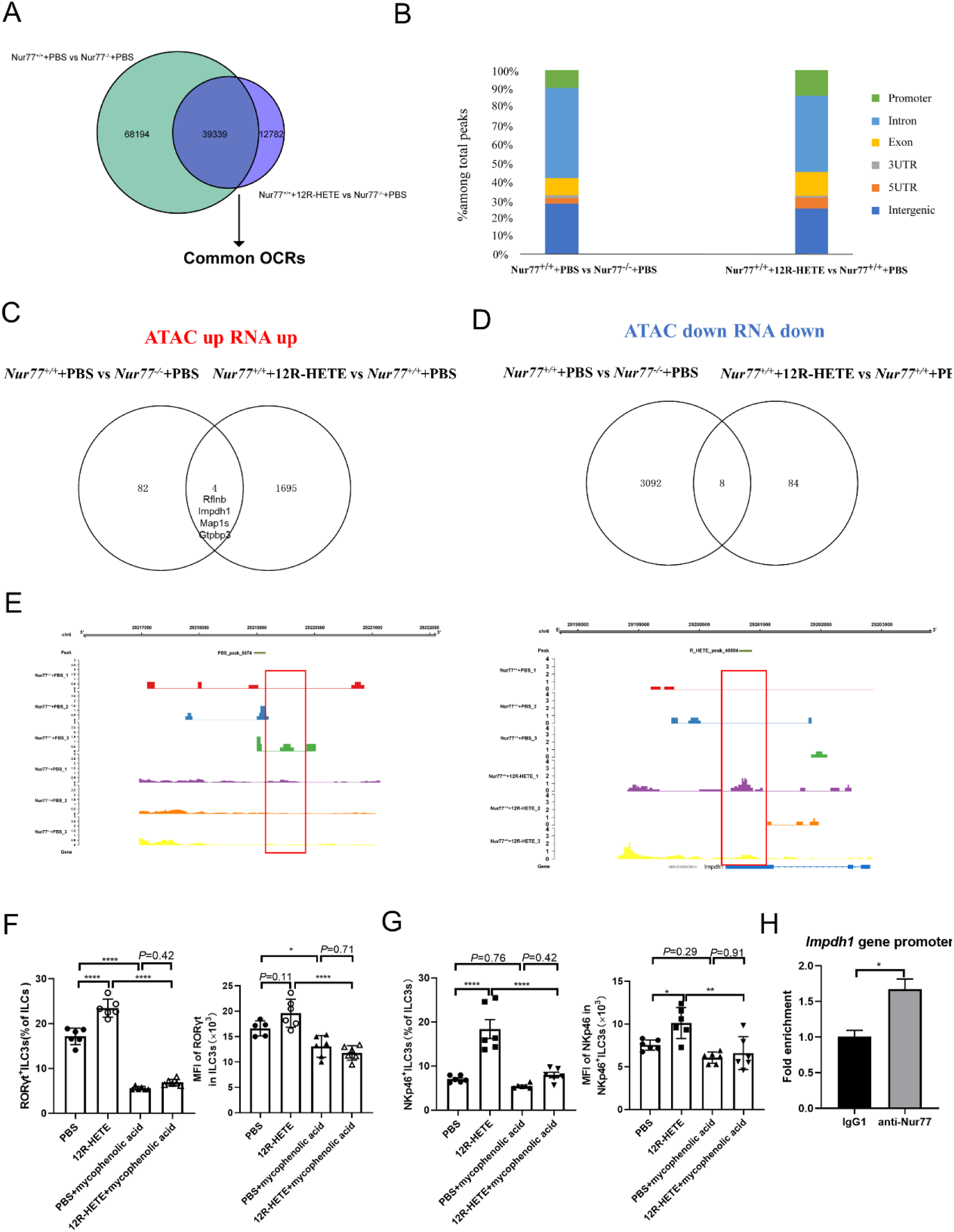
12R-HETE accelerates NKp46^+^ILC3s development by activating Nur77 target genes. ILC3s were sorted from the SI LPLs of mice in *Nur77^+/+^*+PBS group, *Nur77^-/-^*+PBS group and *Nur77^+/+^*+12R-HETE group, respectively. Venn gram depicting numbers of common OCRs in ILC3s after Nur77 knockout or activation (A). Proportions of different peak types among total peaks were shown (B). Venn diagram showed that OCRs related genes were overlapped the up-regulated genes (C) or down-regulated genes (D) from Smart RNA-seq analysis in this study. The percentage of the up-regulated or down-regulated genes in the total overlapping genes was calculated, and the up-regulated overlapping genes were listed. Integrated visualization of ATAC-seq peak at Impdh1 (E). SI LPLs were isolated from mice in four groups at 2-week-old mice and analyzed by flow cytometry for ILC3s (F-G). The percentages of RORγt^+^ILC3s in ILCs (left). MFI of RORγt in RORγt^+^ILC3s (right) (F). The percentages of NKp46^+^ILC3s in ILC3s (left). MFI of NKp46 in NKp46^+^ILC3s (right) (G). Cleavage under targets and release using nuclease (CUT&RUN)-qPCR showing the binding of Nur77 to the promoter region of mouse *Impdh1.* Data are means ± SEM, n = 6. \**P* < 0.05, \*\**P* < 0.01, \*\*\**P* < 0.001.

An integrated analysis of ATAC-seq and Smart RNA-seq showed that there were 4 genes, *Rflnb*, *Impdh1*, *Map1s*, and *Gtpbp3*, which were downregulated and had decreased chromatin accessibility following Nur77 knockout, but were upregulated and had increased chromatin accessibility after 12R-HETE treatment (Fig. 7C). Additionally, 8 genes were upregulated and had enhanced chromatin accessibility after Nur77 knockout, but were downregulated and had decreased chromatin accessibility after 12R-HETE treatment (Fig. 7D). Of the genes upregulated by 12R-HETE, *Impdh1* encodes a protein that is involved in purine nucleotide synthesis and plays a critical role in cell growth and proliferation (Carr et al 1993, Yalowitz and Jayaram 2000), and the ATAC-seq data revealed open chromatin regions in its promoter (Fig. 7E). To investigate whether Impdh1 regulates the percentages and numbers of NKp46^+^ ILC3s by 12R-HETE, a study was carried out in 5-day-old C57BL/6 mice treated with PBS alone, mycophenolic acid, 12R-HETE, or both. The study results showed that administration of 12R-HETE significantly increased the proportion of RORγt^+^ ILC3s and NKp46^+^ ILC3s, but this effect diminished in the presence of mycophenolic acid, suggesting that Impdh1 is required for 12R-HETE to regulate the quantity and ratio of NKp46^+^ ILC3s (Fig. 7F, 7G). We predicted the presence of Nur77 binding sites in the *Impdh1* promoter region by Jaspar. Furthermore, we verified the binding of Nur77 to this site using the technique of cleavage under targets and release using nuclease (CUT&RUN) (Fig. 7H). These results show that Nur77 may play a role in regulating *Impdh1* gene tarascription.

### DISCUSSION

The nuclear receptor Nur77 is an essential player in both innate and acquired immunity in the gut. Previous studies have reported that elimination of *Nur77* exacerbates DSS-induced colitis, while its overexpression by intestinal macrophages and epithelial cells suppresses the production of proinflammatory cytokines (Hamers et al 2015). Also, Nur77 induces apoptosis in T cells and regulates the acquired immune response. Furthermore, Nur77 promotes the expansion of intestinal ILC3s populations by promoting their proliferation (Liu et al 2021). This study revealed a new role of Nur77 in regulating the development of ILC3s, where it modulates their plasticity and boosts their resistance to pathogens, thereby preserving intestinal homeostasis. As a result, this study presents a novel idea of utilizing Nur77 as a target to regulate intestinal immune function and development.

The current research has revealed that Nur77 enhances the proliferation of NKp46^-^ ILC3s while leaving the proliferation of NKp46^+^ ILC3s unaffected. However, it significantly increases the expressions of key transcription factors involved in regulating the differentiation of DN ILC3s to NKp46^+^ ILC3s. Hence, this suggests that Nur77 promotes the expansion of both NKp46^-^ ILC3s and the differentiation of NKp46^-^ ILC3s into NKp46^+^ ILC3s. This allows the quantitative expansion of NKp46^+^ ILC3s without altering their own proliferation. Previous reports have illuminated the role of nutrients, such as vitamin A, vitamin D, arginine, dietary fiber metabolites like short-chain fatty acid, aromatic amino acid metabolites, and ARA metabolite PEG_2_, in regulating the numerical expansion or cellular function of intestinal ILC3s (Kiss et al 2011, van de Pavert et al 2014, Duffin et al 2016, Konya et al 2018, Chun et al 2019, Lin et al 2019). However, there hasn’t been any information on how nutrients regulate intestinal ILC3s plasticity. Considering that 12R-HETE is an ARA metabolite (Powell and Rokach 2015), this research presents critical hints on determining the role of nutrient metabolism in governing intestinal ILC3s plasticity.

To understand Nur77 regulation of ILC3s plasticity at the transcriptional level, Smart RNA-seq, and ATAC-seq technologies were applied to screen effector target genes in this research. The screening process obtained four candidate target genes. Further experimentation with Impdh1 inhibitor revealed its potential as a key target gene mediating the differentiation of Nur77-regulated ILC3s plasticity, linking NKp46^-^ ILC3s to NKp46^+^ ILC3s. Revealing this regulatory mechanism provides a crucial theoretical foundation for comprehending the Nur77 regulation of intestinal ILC3s development. Impdh1 is responsible for encoding the rate-limiting enzyme for guanylate (GMP) synthesis. GMP synthesis provides a basis for rapid cell proliferation, while further metabolism produces guanosine triphosphate (GTP) used for energy in the protein synthesis (Duong-Ly and Kuo 2018). In 3T3L1 cells, Impdh1 was confirmed to be involved in the regulation of adipocyte differentiation (Ruan et al 2002). The results from this study suggest that *Impdh1* may be a key gene in regulating the transition from NKp46^-^ ILC3s to NKp46^+^ ILC3s. This revelation could expand current understanding of the regulatory mechanisms of ILC3s plasticity. It will be interesting to understand whether Impdh1 regulates the conversion of NKp46^-^ ILC3s to Nkp46^+^ ILC3s through signals known to be involved in the regulation of ILC3s plasticity, such as Notch and TGF-β (Chea et al 2016, Viant et al 2016), or through other unknown signaling pathways. In particular, whether other candidates of the targeted genes, such as *Gtbpb3*, also mediate the regulation of ILC3s plasticity by Nur77 requires further confirmation. In addition, Nur77 may also exert regulatory functions through protein/protein interactions. Whether other roles are involved in Nur77 regulation of ILC3s plasticity deserves further investigation.

Nur77 does not have an identified endogenous ligand (Hsu et al 2004). However, recent studies have suggested that ARA, DHA, and PGA_2_ are potential endogenous ligands for Nur77 (Vinayavekhin and Saghatelian 2011, Lakshmi et al 2019). In this study, we conducted a broader survey of lipid metabolites and discovered that 12-HETE is the most abundant ARA metabolite that binds to Nur77-LBD through PMI technology. The binding capacity of 12-HETE was much higher than that of ARA, DHA and PGA_2_ as evidenced by the improved binding relative to the blank control. 12-HETE exists as both 12S-HETE and 12R-HETE in the intestine (Bennett et al 1981, Fretland and Djuric 1989), but analysis using SPR revealed that 12R-HETE has a stronger binding capacity to Nur77 than 12S-HETE and the known exogenous natural agonist Csn-B. Moreover, 12R-HETE demonstrates a stronger activation of the transcriptional regulatory activity of Nur77 than 12S-HETE. 12R-HETE can regulate intestinal ILC3s expansion and plasticity in a Nur77-dependent manner. In contrast, 12S-HETE lacks this ability. Other studies have also recognized significant differences in the biological functions of 12R-HETE and 12S-HETE (Fretland et al 1989, Okuno and Yokomizo 2019). Hence, our findings suggest that 12R-HETE may be the primary endogenous ligand of Nur77 in the intestine.

In conclusion, this study identified 12R-HETE as an endogenous ligand of Nur77 which can bind to Nur77-LBD and activate Nur77 transcriptional activity. 12R-HETE regulates the plasticity of intestinal NKp46^-^ ILC3s toward the NKp46^+^ ILC3s in a Nur77-dependent manner. The Nur77 target gene, *Impdh1*, plays an important role in this process. Overall, these findings advance the understanding of influence of Nur77 on regulating intestinal innate immunity and provide a strong rationale for modulating Nur77 to promote intestinal health.

### METHODS AND MATERIALS

#### Mice

2 weeks old C57BL/6 mice (*Nur77^+/+^*) obtained from the Animal Experiment Center at Huazhong Agricultural University (Wuhan, China) were used for the current study. *Nur77^-/-^* mice were obtained from Cyagen US Inc (Guangzhou, China). Mice were housed under specific pathogen-free conditions in an air-conditioned room at 22 ± 4 ℃ with a 12 h light/dark cycle. Food and water were supplied ad libitum. All animal experimental protocols were approved by the Institutional Animal Care and Use Committee of Huazhong Agricultural University.

#### Drug treatment

Neonatal mice at postnatal 4–7 days and postnatal 8–14 days were separately provided 50 or 100 μL PBS, 1 μmol 12S-HETE or 1 μmol 12R-HETE treated for 10 d. In the experiments of mechanism, mice were intraperitoneally injected with Notch inhibitors or Impdh1 inhibitors after treating with PBS or 12R-HETE, mice were dissected after 10 d.

#### Salmonella Typhimurium infection model

The *Salmonella Typhimurium* strain (SL1344) was grown with shaking overnight at 37 °C in Luria Bertani (LB) broth subcultured for 4 h, and washed twice with cold PBS prior to use. For oral *Salmonella Typhimurium* infections, mice were gavaged orally with OD600 = 0.8 ∼ 1.0 of bacteria in 50 μL PBS and corresponded to 10^8^ bacteria per mL as confirmed by plating of the inoculum. Mice were weighed daily, then the mice were dissected on day 1 and day 3 after infections. The small intestines were collected to isolate intestinal lymphocyte cells. The tissues of liver and spleen were collected at different time points and homogenized in 1 mL of sterile PBS, serial dilutions of the homogenates were plated on Bismuth Sulfite Agar Medium, 37 °C cultured for 12 h to determine the number of *S. typhimurium*.

#### Histological analysis

Colons were cleaned with PBS and were fixed with 4 % paraformaldehyde. Tissues were then embedded in paraffin, sectioned and stained with hematoxylin and eosin. The following histological parameters were evaluated: leukocyte infiltration, epithelial injury and submucosal oedema.

#### Mouse small intestinal Lipids extraction

The lipid extraction protocol used here was adopted from Folch (Folch et al 1957). Briefly, the mouse small intestinal lipids of 2 week-old mice were extracted with 10 mL chloroform/methanol mixture (2: 1, v /v), crude extracts were washed by 2 mL water or salt solution, the two liquids are mixed with a stirring rod and centrifugated 20 minutes at 2400 rpm. The supernatant was collected and then evaporated to dryness under nitrogen gas stream, reconstituted in 100 μL of methanol: water (1:1, v/v).

#### LC-MS/MS analyzed protein–metabolite interactions

The identify potential small-molecule ligands for Nur77 using protein metabolite interactions was done as previously described (Qin et al 2019). There are two experimental groups in our study. 150 μL of 500 μg/mL GST-Nur77-LBD was incubated with the resins. For control, GST-Nur77-LBD without lipids extraction at 4 ℃ for 3 h with shaking. The lipids extracted from mouse small intestine were incubated with GST-Nur77-LBD at 4 ℃ for 3 h with shaking. The resins were washed by PBS buffer containing 8% DMSO/EtOH and 50 mM imidazole for three times. The protein bound lipids were eluted by incubating the resin with of PBS buffer containing 500 mM imidazole for 30 min methanol was added to the elution solution to precipitate protein. Subsequently, the supernatant was lyophilized, and it was analyzed by LC-MS/MS after redissolved in methanol: water (1:1, v/v).

LC-MS/MS was carried out Metware (Wuhan, China) by using an LC-ESI-MS/MS system (HPLC, Shim-pack UFLC SHIMADZU CBM30A system; MS, Applied Biosystems 6500 Q TRAP system). The condition of LC-MS/MS was done as previously described (Huang et al 2022).

#### Construction of plasmid

The four NBFI-B response elements (NBRE) (AAAGGTCA) were inserted into the pGL3-promoter vector (Promega). GAL4-DBD was inserted into the pGL3-promoter vector (Promega). Nur77-LBD was inserted into the pcDNA3.1 vector (Promega).

#### Luciferase reporter assays

The NBRE luciferase vector and renilla luciferase-expressing plasmid (pTK) were transfected into HEK293T cells for 6 h. Then, these cells were treated with Csn-B, 12S-HETE, and 12R-HETE for 14 h. After washing with PBS, cells were lysed using Dual-Glo luciferase Dual-Luciferase® Reporter Assay system (Promega). The luciferase values were normalized to the Renilla values. The transfection experiments were performed in triplicate for each independent experiment.

#### Surface plasmon resonance

To analyze the binding of Nur77 and Csn-B, 12S-HETE, 12R-HETE, Nur77-LBD protein was fixed on the COOH sensor chip by capture-coupling. In this study, Csn-B act as positive control, 12S-HETE at the concentrations of 1.5, 3, 6.25 mmol/L and 12R-HETE at the concentrations of 5, 10, 20, 40, 80 mmol/L were injected sequentially into the chamber in PBS running buffer, and the interactions of Nur77-LBD with Csn-B, 12S-HETE and12R-HETE were detected by Open SPRTM (Nicoya Lifesciences, Waterloo, Canada) at 25 °C. The binding time and disassociation time were both 250 seconds; the flow rate was 20 μL/min, and the chip was regenerated with 0.25 % SDS. A one-to-one diffusion-corrected model was fitted to the wavelength shifts corresponding to the varied drug concentrations. The data were retrieved and analyzed with Trace Drawer software (Ridgeview Instruments AB, Sweden).

#### Molecular docking

The human Nur77 ligand-binding domain (LBD) was obtained from the Protein Data Bank (PDB code 3V3Q). Human Nur77 was used to predict the structure of porcine Nur77 and define the active site of porcine Nur77 by Swiss model website. A square three-dimensional structure with the size of 10×10×10 ^A^ was established to form grid points for molecular docking. The compounds were prepared in the range of pH5∼9, conformational search was conducted, and 3D structures were obtained for docking. Connect with extra precision (XP) and flexible mode.

#### Isolation of small Intestinal lamina propria lymphocytes

Isolation of lamina propria cells were performed as follows. Small intestines were dissected and fat tissues were removed. Intestines were dissected longitudinally and washed with cold PBS. Intestines were then cut into 1 cm pieces. Intestines were incubated in PBS containing 1mmol dithiothreitol (DTT) for 10 min at room temperature (RT) and 30mM Ethylenediaminetetraacetic acid (EDTA) at 37 °C for 10 min for two cycles. Intestines washed with PBS, then digested in RPMI1640 medium (Gibico) containing DNase I(Sigma) (150 μg/mL) and collagenase VIII (Sigma) (100 U/mL) at 37 °C in 5 % CO_2_ incubator for1.5 h. The digested tissues were homogenized by vigorous shaking and passed through 70 μm cell strainer. Mononuclear cells were then collected from the interphase of 40/80% Percoll gradients (GE Healthcare Life Sciences) after spinning at 400g for 10 min at RT.

#### Flow cytometry and sorting of ILC3s

Dead Cells were detected with Fixable Viability Stain 510 (BD Pharmingen). Mouse CD16/32 antibody (2.4G2, BD Pharmingen) was used to block the non-specific binding to Fc receptors before all surface staining. All antibodies were purchased from BD Pharmingen unless otherwise specified. For surface marker staining, we used antibodies to the following mouse proteins: CD45 (30-F11), lineage markers (CD3e/CD11b/CD45R/B220Ly-76/ Ly-6G/ Ly-6C), CD127(SB/199). For detection of intracellular cytokine production, cells were stimulated with phorbol-12-myristate13-acetate (PMA) (50 ng/mL) and ionomycin (500 ng/mL, Sigma-Aldrich) for 2h, and Brefeldin (2 μg/mL, BioLegend) for 2 h. Cells were subsequently surface-stained with a combination of the antibodies listed above, fixed and permeabilized using Transcription Factor Buffer Set and stained with IL-17A-PerCP-Cyanine5.5 (eBio17B7, eBioscience), IL-22-APC (IL22JOP, eBioscience) and IFN-γ-FITC (XMG1.2). For measurement of transcription factor expression, cells were fixed and permeabilized according to the manufacturer’s instructions of Transcription Factor Buffer Set, and then stained with RORγt (Q31-378). The antibodies used for sorting ILC3s were Anti-Mouse CD45 (30-F11) (BD Pharmingen), Fixable Viability Stain 510 (BD Pharmingen), PerCP-Cy5.5 Lineage Antibody, FITC Rat Anti-Mouse CD90.2(53-2.1), PE Hamster Anti-Mouse KLRG1(2F1), PE-Cy7 Mouse Anti-Mouse NK-1.1(PK136), APC CD335 (NKp46) Antibody (29A1.4) (Biolegend). Flow cytometry data were collected using cytoflex-LX (Beckman Coulter) and analyzed with FlowJo software. ILC3s were sorted from *Nur77*^+/+^ with PBS, *Nur77*^+/+^ with 12R-HETE and *Nur77*^-/-^ with PBS mice using Moflo XDP (Beckman Coulter)。

#### Cleavage Under Targets and Release Using Nuclease (CUT&RUN) -qPCR

The operation of experiments was followed the instruction of Hyperactive pG-MNase CUT&RUN Assay Kit for qPCR (Vazyme). Briefly, Sorted ILC3s (5×10^4^ cells/reaction, n=3) were incubated and combined with magnetic beads precoated with concanavalin A (Vazyme). Cells were permeabilized and incubated with Chip grade purified anti-Nur77 antibody (10 μg/mL) or purified mouse IgG1 monoclonal-isotype control antibody (10 μg/mL, abclonal) for 2 hours at room temperature. After washing, protein G-MNase was added and incubated on ice for 1 hour. Then Cacl_2_ was added to activate MNase and cleave chromatin. The reaction was stopped by adding stop buffer. DNA fragments was extracted and purified using a FastPure gDNA Mini Kit. The following primer sequence were used to amplify the fragment of the *Impdh1* gene that contains the potential Nur77 binding site: forward, 5’-GGAATTGCAGGAAGAGGGAGGG-3’; reverse, 5’-GGGCTCAAGGCTGGGTTAGG-3’. Spike in DNA was used to normalizes.

#### Smart Sequencing

Total small intestinal ILC3s RNA exacted from the three different treatments was used to construct smart RNA sequencing (sRNA). RNA libraries were sequenced on an Illumina HiSeq platform by Wuhan Igenebook Biotechnology Co.Ltd (Wuhan, China). After obtaining raw data, these were screened to eliminate adaptor sequences in reads. We selected sRNA ranging from 18–30 nt in clean reads obtained from the last step and summarised the types and abundance of sRNA. The distribution of common and unique sRNA between three different groups was also analyzied.

#### ATAC Sequencing

ATAC-seq was performed according to previously described (Wu et al 2016). Briefly, small intestinal ILC3s were lysed in lysis buffer (10 mM Tris-HCl (pH 7.4), 10 mM NaCl, 3 mM MgCl_2_ and NP-40) for 10 min on ice to prepare the nuclei, then, nuclei were pelleted by centrifugation at 500 g at 4 ℃for 10 min to remove the supernatant. The quality of nuclei was checked under microscope by 49, 6-diamidino-2-phenylindole staining. Crude nuclei were incubated with the tagmentation reaction mixture which contained Mix 25 mL of reaction buffer, 2.5 mL of NexteraTn5 transposase, and 22.5 mL of nuclease free H_2_O at 37 °C for 30 min. The Qiagen MinElute PCR purification kit was used immediately to purify DNA after transposition. Sequencing library was constructed using the HiSeq X Ten platform and 150-bppaired-end sequencing by Igenebook Biotechnology Co., Ltd. (Wuhan, China).

#### Statistical analysis

Before data analysis, Normality and Lognormality Tests and Shapiro-Wilk were used for normality tests (significance level was set at 0.05). According to whether the data fit the normal distribution, the unpaired T test or unpaired one-way analysis of variance or Kruskal-Wallis test was used for analysis, respectively. Correlation analysis was performed using GraphPad Prism 8.3.0 software, and the data were presented as mean ± standard error. Ns: not significant. \**P* < 0.05, \*\**P* < 0.01, \*\*\**P* < 0.001.

## ACKNOELEDGEMENTS

We thank Yangene biotechnology Co., Ltd. for the performance of SPR and Molecular docking experiments. Thanking Metware (Wuhan, China) for analysing. We would like to thank Wuhan Igenebook Biotechnology Co.Ltd for Smart RNA seq and ATAC seq high-throughput sequencing.

## COMPETING OF INTERENTS

The authors declare no competing interests.

## AUTHOR CONTRIBUTIONS

NH, HW and JP designed experiments. NH and LY executed experiments and /or performed data analysis. NH and HW writing—original draft preparation. LY and HL performed breeding of mice. HW and JP provided funding support. All authors have read and agreed to the published version of the manuscript.

## ETHICS APPROVAL

The mice experiments were approved by the Institutional Animal Care and Use Committee of Huazhong Agricultural University (HZAUMO-2023-0098).

## DATA AVAILABILITY

The data are included in this article and the supplemental data files

---

**supplementary Fig. 1.**
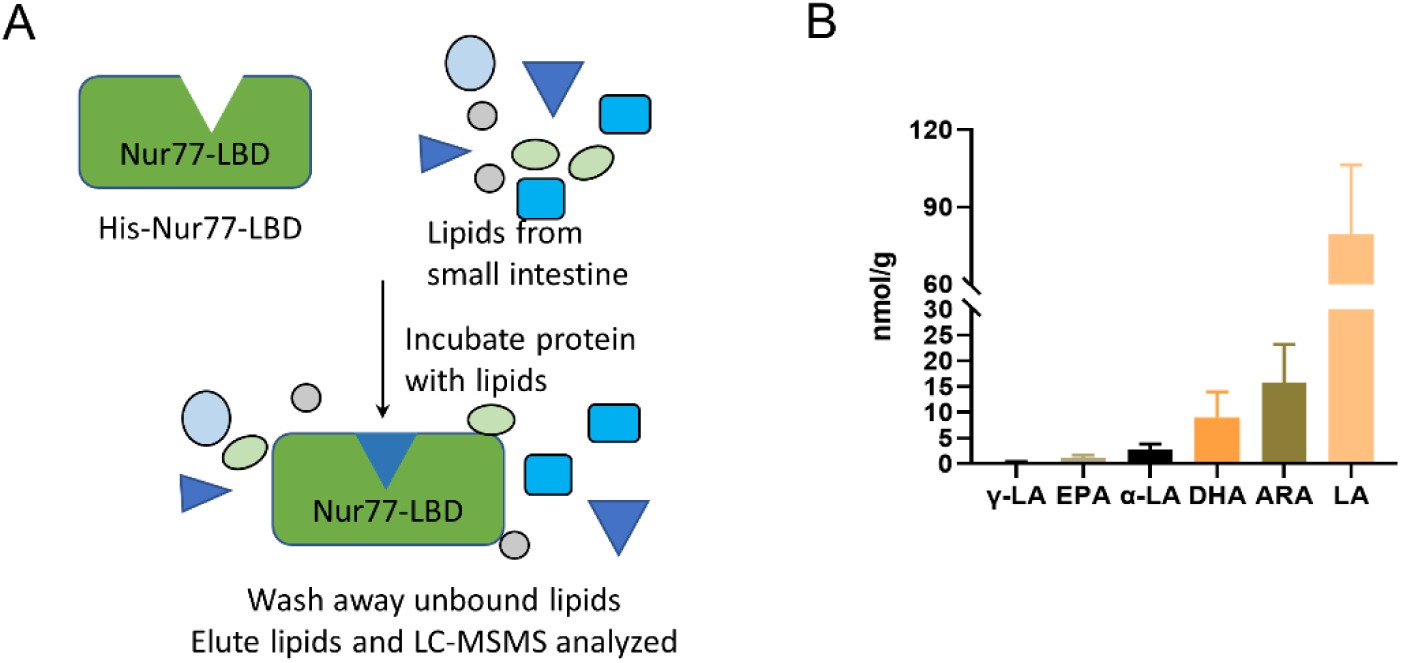
Extraction and identification of ARA metabolites from mice intestine bound to Nur77-LBD. Metabolomics workflow for the identification of PMIs with Nur77LBD (A). Absolute concentration of polyunsaturated fatty acids bound to Nur77-LBD (B).

**supplementary Fig. 2.**
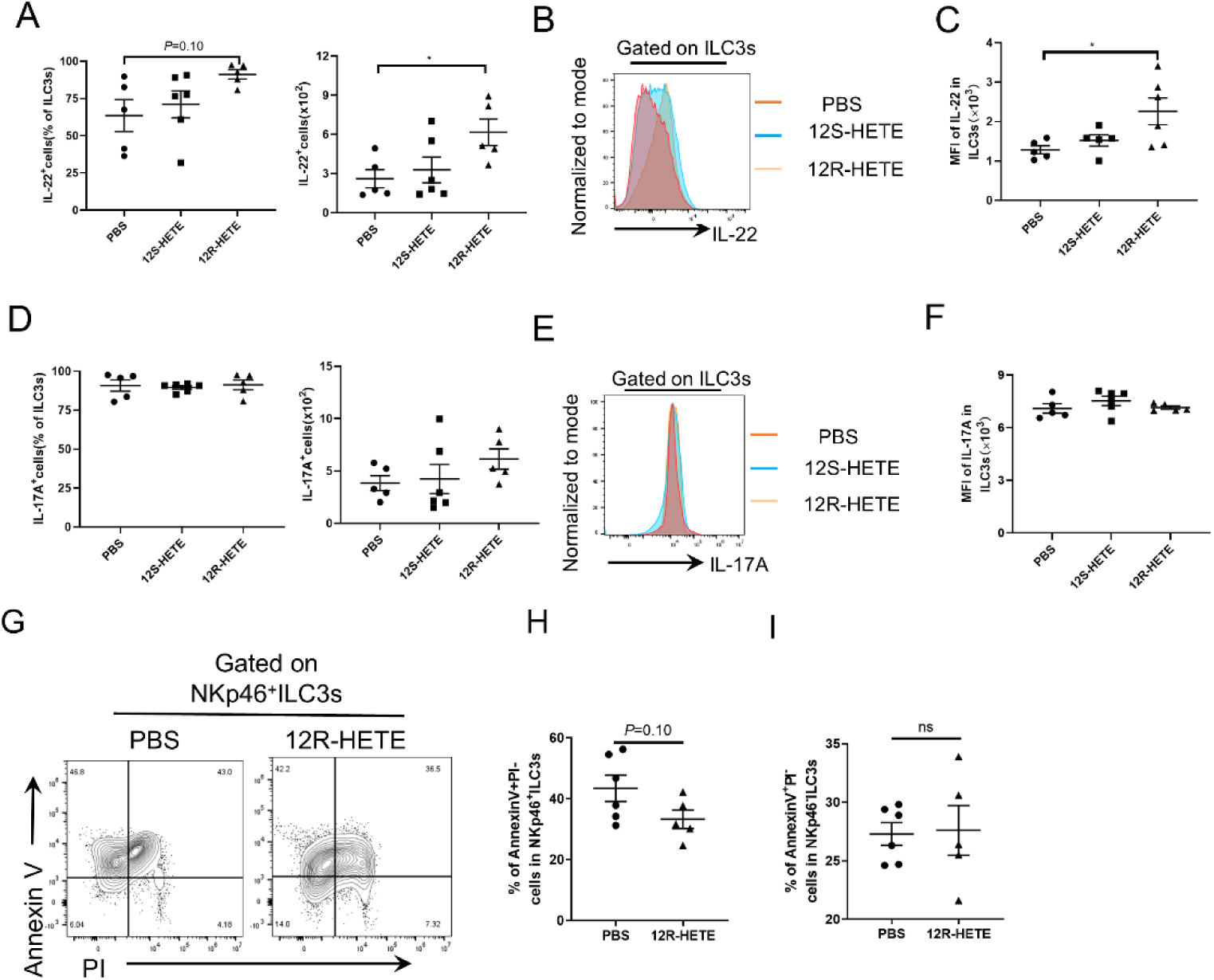
Effects of 12R-HETE on IL-17A, IL-22 and apoptosis produced by ILC3s. Percentages and absolute numbers of IL-22 from ILC3s in ILC3s (A). Representative histograms show the expression of IL-22 in ILC3s. The PBS group is red curves, 12S-HETE group is blue curves, and 12R-HETE is orange curves (B). MFI of IL-22 in ILC3s (C). Percentages (left), absolute numbers (right) and MFI of IL-17A from ILC3s in ILC3s from different treatment groups were shown by flow cytometry (D-F). Numbers in flow plots represent percentages of apoptotic cells on NKp46^+^ ILC3s gate were shown on the left. Percentages of apoptotic cells in NKp46^+^ ILC3s were shown on the right (G). Percentages of apoptotic cells in NKp46^+^ ILC3s (H) or NKp46^-^ ILC3s (I) were shown. Data are mean ± SEM, n=5-6.

**supplementary Fig. 3.**
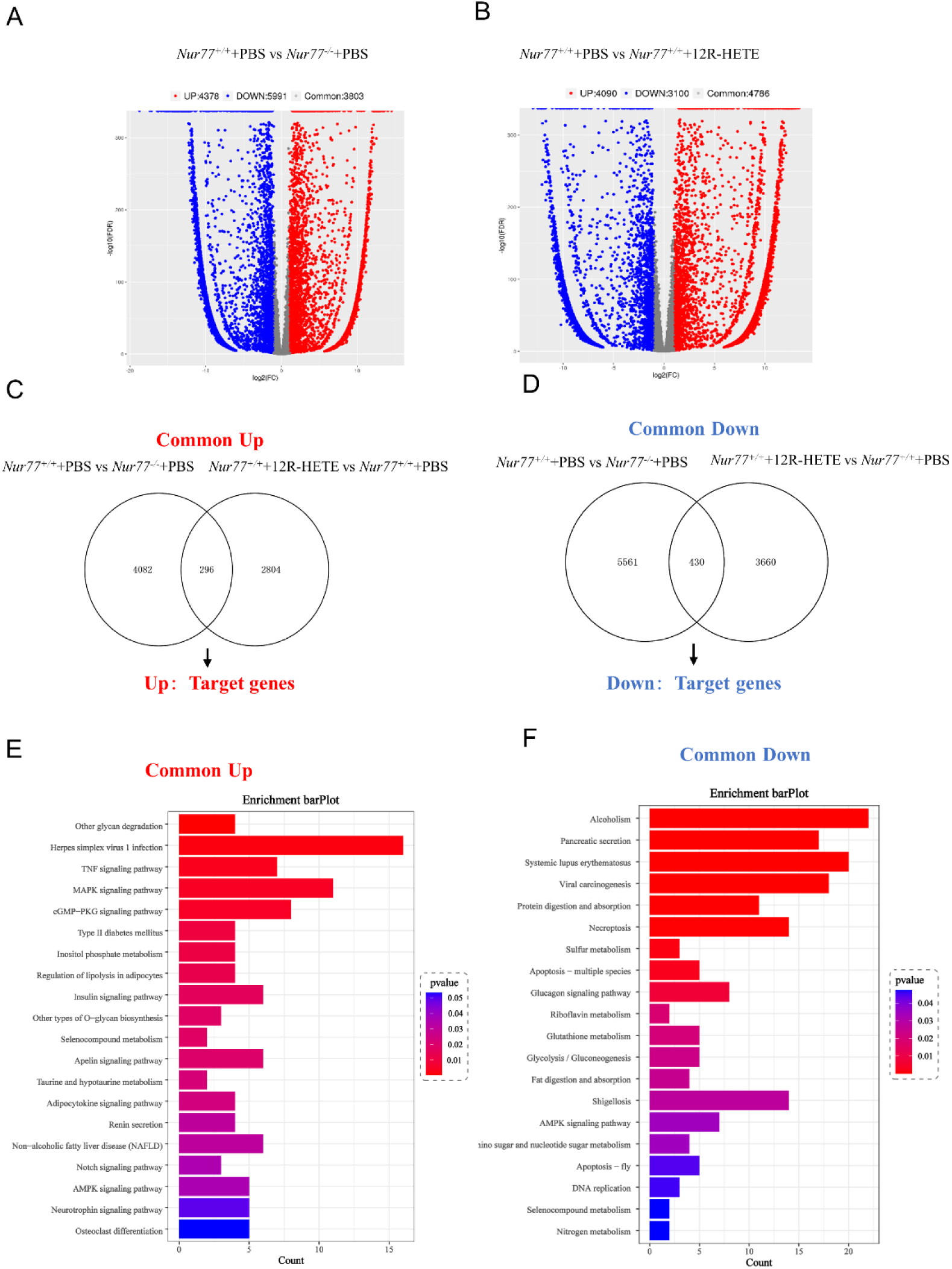
Smart RNA seq and ATAC seq were used to analyze differential genes. ILC3s were sorted from the SI LPLs of mice in *Nur77^+/+^*+PBS group, *Nur77^-/-^*+PBS group and *Nur77^+/+^*+12R-HETE group, respectively. Volcano map shows DEGs detected by Smart RNA seq (A-B). Venn gram depicting numbers of common differential genes in ILC3s cells after Nur77 knockout or activation (C-D). KEGG analysis of the co-upregulated or co-downregulated DEGs after Nur77 knockout or activation (E-F).

