## Supplemental Data 1 for "12R-HETE acts as an endogenous ligand of Nur77 in intestine and regulates ILC3s plasticity"

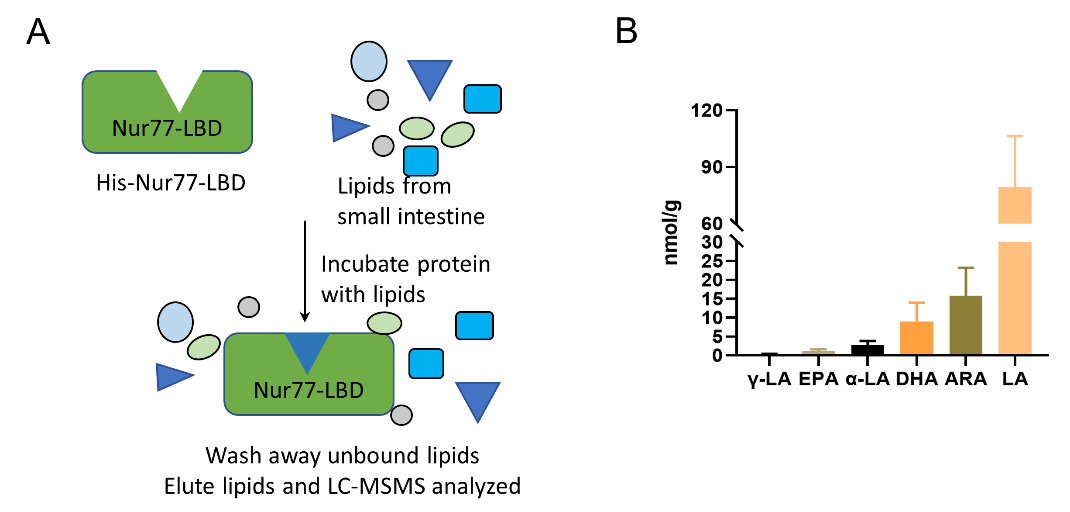

**supplementary Fig. 1 Extraction and identification of ARA metabolites from mice intestine bound to Nur77-LBD**

Metabolomics workflow for the identification of PMIs with Nur77LBD (A). Absolute concentration of polyunsaturated fatty acids bound to Nur77-LBD (B).

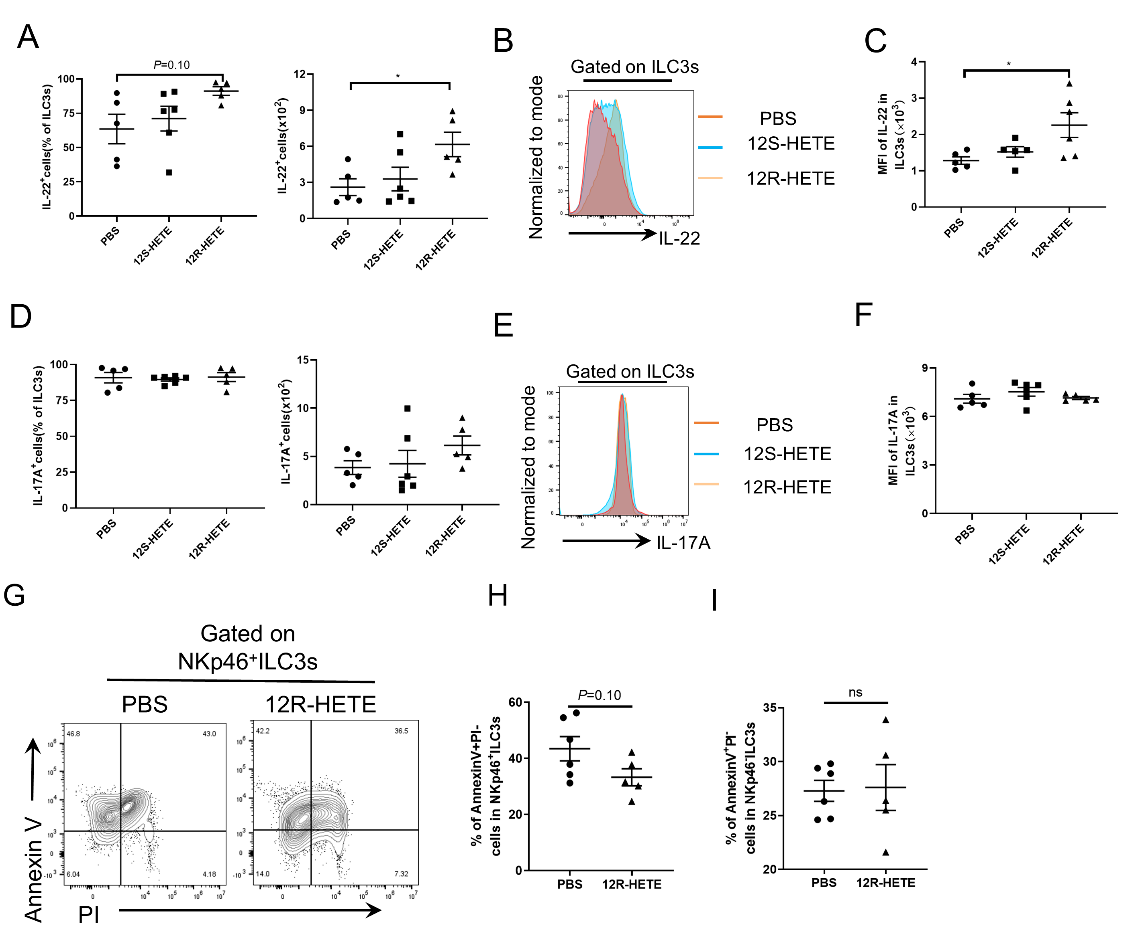

**supplementary Fig. 2 Effects of 12R-HETE on IL-17A, IL-22 and apoptosis produced by ILC3s**

Percentages and absolute numbers of IL-22 from ILC3s in ILC3s (A). Representative histograms show the expression of IL-22 in ILC3s. The PBS group is red curves, 12S-HETE group is blue curves, and 12R-HETE is orange curves (B). MFI of IL-22 in ILC3s (C). Percentages (left), absolute numbers (right) and MFI of IL-17A from ILC3s in ILC3s from different treatment groups were shown by flow cytometry (D-F). Numbers in flow plots represent percentages of apoptotic cells on NKp46^+^ ILC3s gate were shown on the left. Percentages of apoptotic cells in NKp46^+^ ILC3s were shown on the right (G). Percentages of apoptotic cells in NKp46^+^ ILC3s (H) or NKp46^-^ ILC3s (I) were shown. Data are mean ± SEM, n=5-6.

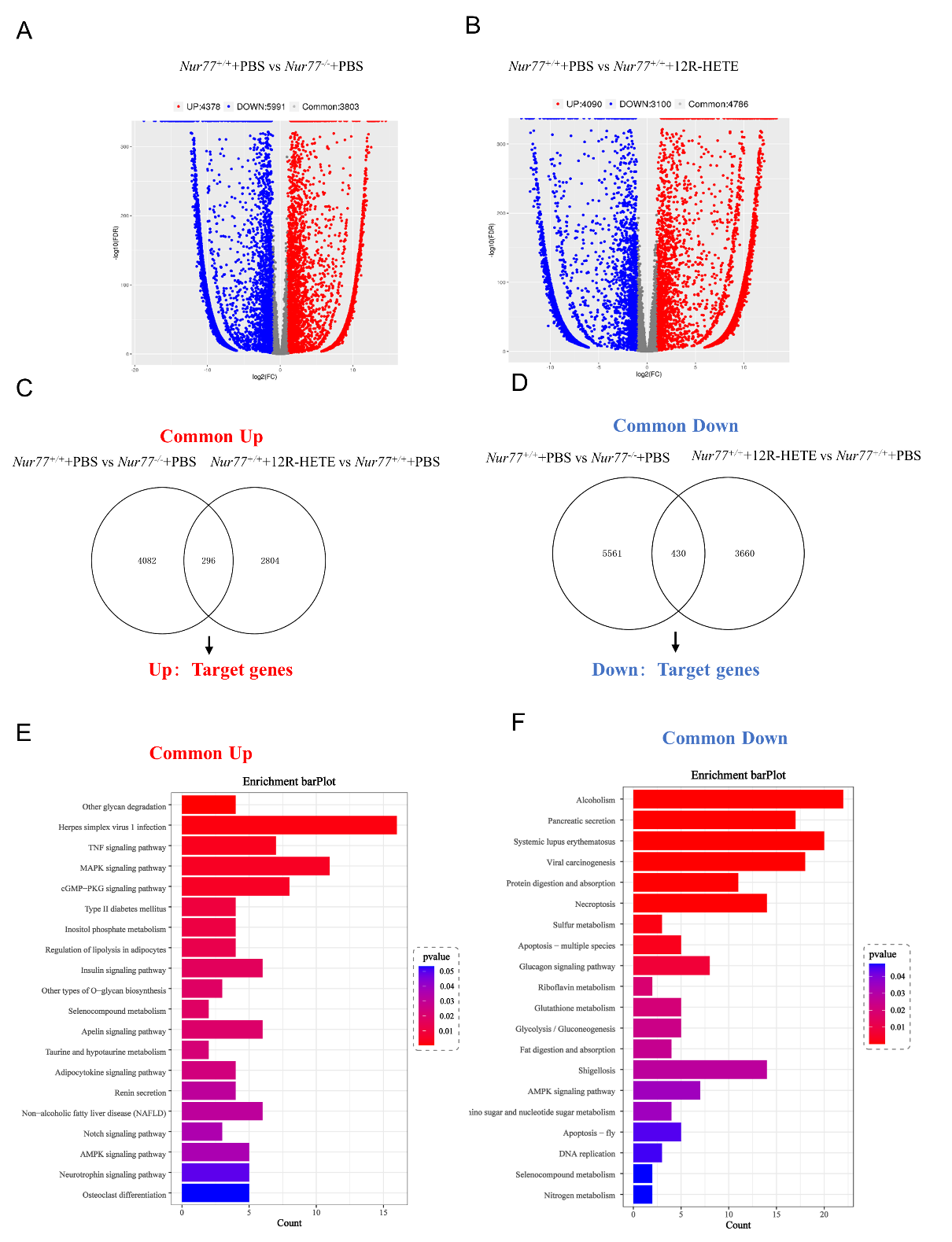

**supplementary Fig. 3 Smart RNA seq and ATAC seq were used to analyze differential genes**

ILC3s were sorted from the SI LPLs of mice in *Nur77^+/+^*+PBS group, *Nur77^-/-^*+PBS group and *Nur77^+/+^*+12R-HETE group, respectively. Volcano map shows DEGs detected by Smart RNA seq (A-B). Venn gram depicting numbers of common differential genes in ILC3s cells after Nur77 knockout or activation (C-D). KEGG analysis of the co-upregulated or co-downregulated DEGs after Nur77 knockout or activation (E-F).
